# CircularLogo: A lightweight web application to visualize intra-motif dependencies

**DOI:** 10.1101/098327

**Authors:** Zhenqing Ye, Tao Ma, Michael T. Kalmbach, Surendra Dasari, Jean-Pierre A. Kocher, Liguo Wang

## Abstract

**Background:** The sequence logo has been widely used to represent DNA or RNA motifs for more than three decades. Despite its intelligibility and intuitiveness, the traditional sequence logo is unable to display the intra-motif dependencies and therefore is insufficient to fully characterize nucleotide motifs. Many methods have been developed to quantify the intra-motif dependencies, but fewer tools are available for visualization.

**Result:** We developed *CircularLogo*, a web-based interactive application, which is able to not only visualize the position-specific nucleotide consensus and diversity but also display the intra-motif dependencies. Applying *CircularLogo* to HNF6 binding sites and tRNA sequences demonstrated its ability to show intra-motif dependencies and intuitively reveal biomolecular structure. *CircularLogo* is implemented in JavaScript and Python based on the Django web framework. The program’s source code and user’s manual are freely available at http://circularlogo.sourceforge.net. *CircularLogo* web server can be accessed from http://bioinformaticstools.mayo.edu/circularlogo/index.html.

**Conclusion:** *CircularLogo* is an innovative web application that is specifically designed to visualize and interactively explore intra-motif dependencies.

## Background

Many DNA and RNA binding proteins recognize their binding sites through specific nucleotide patterns called motifs. Motif sites bound by the same protein do not necessarily have same sequence but typically share consensus sequence patterns. Several methods have been developed to statistically model the position-specific consensus and diversity of nucleotide motifs using the position weight matrix (PWM) or position-specific scoring matrix (PSSM) [1,2]. These mathematical representations are usually visualized using sequence logos, which depict the consensus and diversity of each motif residue as a stack of nucleotide symbols. The height of each symbol within the stack indicates its relative frequency, and the total height of symbols is scaled to the information content of that position [3,4].

Traditional PWM and PSSM assume statistical independence between nucleotides of a motif. However, such assumption is not completely justified, and accumulated evidence indicates the existence of intra-motif dependencies [5-8]. For example, an analysis of wild-type and mutant *Zif268* (*EGR-1*) zinc fingers, using microarray binding experiments, suggested that the nucleotides within transcription factor binding site (TFBS) should not be treated independently [5]. In addition, the intra-dependences within a motif were also revealed by a comprehensive experiment to examine the binding specificities of 104 distinct DNA binding proteins in mouse [8]. Intra-motif dependencies when into consideration could substantially improve the accuracy of *de novo* motif discovery [9]. Therefore, many statistical methods have been developed to characterize the intra-motif dependencies, which include the generalized weight matrix model [10], sparse local inhomogeneous mixture model (Slim) [11], transcription factor flexible model based on hidden Markov models (TFFMs) [12], the binding energy model (BEM) [13], and the inhomogeneous parsimonious Markov model (PMM) [14]. However, the most commonly used visualization tools such as WebLogo [3] and Seq2Logo [15] are incapable of displaying these intra-motif dependencies.

Only a handful of tools like CorreLogo, enoLOGOS, and ELRM are capable of visualizing positional dependencies [16,17,18]. CorreLogo depicts mutual information from DNA or RNA alignment using three-dimensional sequence logos generated via VRML and JVX. However, CorreLogo’s three-dimensional graphs are difficult to interpret because of the excessively complex and distorted perspective associated with the third dimension. ELRM generates static graphs to visualize intra-motif dependences. ELRM splits up “base features” and “association features” and fails to comprehensively integrate nucleotide diversities and dependencies. In addition, ELRM is limited to measuring dependence with its own built-in method. Similar to ELRM, enoLOGOS represents the dependency between different positions using a matrix plot underneath the nucleotide logo. While pLogo allows user to visualize correlations to a particular nucleotide position, it fails to provide overall view of intra-motif dependencies [4]. Finally, all of these tools lack the functionality for users to explore and interpret the data in an interactive fashion.

In this study, we developed *CircularLogo*, an interactive web application, which is capable of simultaneously displaying position-specific nucleotide frequencies and intra-motif dependencies. *CircularLogo* uses an open-standard, human-readable, flexible and programming language independent JSON (JavaScript Object Notation) data format to describe various properties of DNA motifs. Other commonly used motif formats such as MEME, TRANSFAC, and JASPAR can be easily converted into JSON format.

## Implementation

### JSON-Graph specifications of nucleotide motif representation

We used the JSON-Graph format to describe nucleotide motif in order to make it intelligible and malleable. The schema of JSON-Graph format is illustrated as below:

~~~
{
"id": "Toy motif",
"background":{"key":["A","T","C","G"],"val":[0.25,0.25,0.25,0.25]},
"pseudocounts":{"key":["A","T","C","G"],"val":[0.25,0.25,0.25,0.25]},
"nodes": [
           {"index": 0, "label": "1", "bit": 0.78, "base": ["A", "T", "C", "G"], "freq": [0.006, 0.043, 0.347, 0.604]},
           {"index": 1, "label": "2", "bit": 1.57, "base": ["T", "C", "A", "G"], "freq": [0.012, 0.017, 0.032, 0.939]},
           {"index": 2, "label": "3", "bit": 0.61, "base": ["C", "G", "A", "T"], "freq": [0.027, 0.053, 0.388, 0.532]},
           ……
],
"links": [
           {"source": 0, "target": 1, "value": 2.0},
           {"source": 2, "target": 4, "value": 8.0},
           {"source": 2, "target": 7, "value": 6.0},
           ……
]
}
~~~

The contents within two curly braces describe a DNA or RNA motif. Specifically, the “*id*” keyword specifies the name of the motif. The “*background*” keyword designates nucleotides frequencies (in the order of A, T, C and G) of the relevant genomic background. For example, when studying motifs in human genome, these percentages are computed from the human reference genome as background distribution. By default, they are set to 0.25 representing equal frequencies. The “*pseudocounts*” keyword represents the extra nucleotides added to each position of the motif to avoid zero-division error in small data set; these are set to 0.25 for each nucleotide by default. The “*nodes*” section describes various properties of motif residues using the following keywords: a) the “*index*” keyword specifies the sequential order (in anticlockwise) of nucleotide stacks b) the “*label*” keyword denotes the identity of each nucleotide stack c) the “*bit*” keyword refers to the information content calculated for each nucleotide stack d) the “*base*” keyword indicates the four nucleotides sorted incrementally by their corresponding frequencies as designated by the “*freq*” keyword. The “*links*” section describes the pairwise dependencies between nucleotide stacks using the following keywords: a) the “*source*” and “*target*” keywords denoting the start and the end positions of nucleotide stacks b) the “value” keyword indicates the width of the link that is proportional to the strength of dependence between the two linked positions.

### *CircularLogo* web server

*CircularLogo* web application uses NGINX (https://www.nginx.com/) web server with uWSGI (https://pypi.python.org/pypi/uWSGI) gateway interface to handle multiple concurrent client requests. The application is hosted on Amazon Elastic Compute Cloud (Amazon EC2).

### Measure intra-motif dependencies using χ^2^ statistic

We implemented two metrics to calculate the dependence between a pair of nucleotide positions: mutual information and the χ^2^ statistic. The χ^2^ statistic is widely used to test the independence of two categorical variables and corresponding *Q* score is a natural measure of dependency between two events that quantifies the co-incidence as follows. Let us assume that a DNA motif is *l* nucleotides long and is built from *N* sequences. For given two positions *i* and *j* within the motif (1≤ *i* ≤ *l*, 1≤ *j* ≤ *l*, *i* ≠ *j*), the observed dinucleotide frequency is denoted as *O_ij_*, which can be obtained by counting di-nucleotide combinations from the input *N* sequences. The expected di-nucleotide frequency is represented as *E_ij_*. The χ^2^ statistic score is then calculated as:

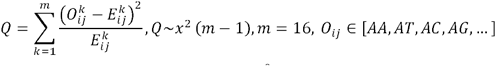

Here, *m* is the total number of di-nucleotides (4^2^ = 16).

### Measure intra-motif dependencies using mutual information

The second built-in approach to measure dependence is the mutual information. This metric quantifies the mutual dependence between two discrete random variables *X* (X = [*A,C,G,T*]) and *Y* (Y = [*A,C,G,T*]) and it is defined as:

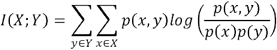

Here, *x* (*x* ∈ [*A,C,G,T*]) and *y* (*y* ∈ [*A,C,G,T*]) represent nucleotides at two nucleotide stacks X and Y, respectively. *p*(*x*) and *p*(*y*) denote the nucleotide frequencies of *x* and *y*. *p*(*x*, *y*) defines the frequencies of dinucleotides (*xy*) from *X* and *Y*.

### HNF6 motif analysis

HNF6 ChIP-exo data was obtained from Array Express (accession number E-MTAB-2060; http://www.ebi.ac.uk/arrayexpress/experiments/E-MTAB-2060/), processed with MACE[19], and HNF6 binding sites were extracted. The 5549 65-nucleotide (upstream 20 nucleotides + 25 nucleotides HNF6 binding site + downstream 20 nucleotides) sequences were published to https://sourceforge.net/projects/circularlogo/files/test/. All sequences were aligned by the HNF6 motif, which start from postion-29 to position-36.

### tRNA sequence analysis

A total of 1114 tRNA sequences were downloaded from RFAM database [23] in the form of RFAM ‘seed’ alignment format (accession # RF00005; https://correlogo.ncifcrf.gov/ccrnp/trnafull.html). After excluding sequences with gaps in the alignment, 291 sequences were used as the final dataset to generate circular logo of tRNA (https://sourceforge.net/projects/circularlogo/files/test/). Mutual information was used as the metric to measure intra-motif dependencies. The lower 33% links were filtered out.

## Results

### Circular nucleotide motif

Unlike the traditional sequence logos that display motif residues on a two-dimensional Cartesian coordinate system (with the x-axis denoting the position of residue stacks and the y-axis denoting the information contents), *CircularLogo* visualizes motifs using a polar coordinate system that facilitates the display of pairwise intra-motif dependencies with linked ribbons (Fig. 1). Since traditional PWM or PSSM representations do not preserve intra-motif dependency information, we use the JSON-Graph as the main input format to *CircularLogo*. When the input file is in JSON-Graph format that has pre-calculated nucleotide frequencies and dependencies, the *CircularLogo* simply transforms this file into a pictorial representation. In addition, *CircularLogo* also accepts the FASTA format motif representation as input. In this scenario, *CircularLogo* transforms the FASTA information into a JSON-Graph format by calculating the intra-motif dependency using the built-in χ^2^ statistic or mutual information metric, and determine the height of each nucleotide stack in the same way as webLogo [3]. In brief, *CircularLogo* generates a sector for each motif position and draws nucleotide stack within that sector based on the information content and relative frequencies of nucleotides. All sectors are properly arranged into a circular layout. The width of linked arcs indicates the strength of intra-dependency between each pair of nucleotide positions.

**Figure 1.**
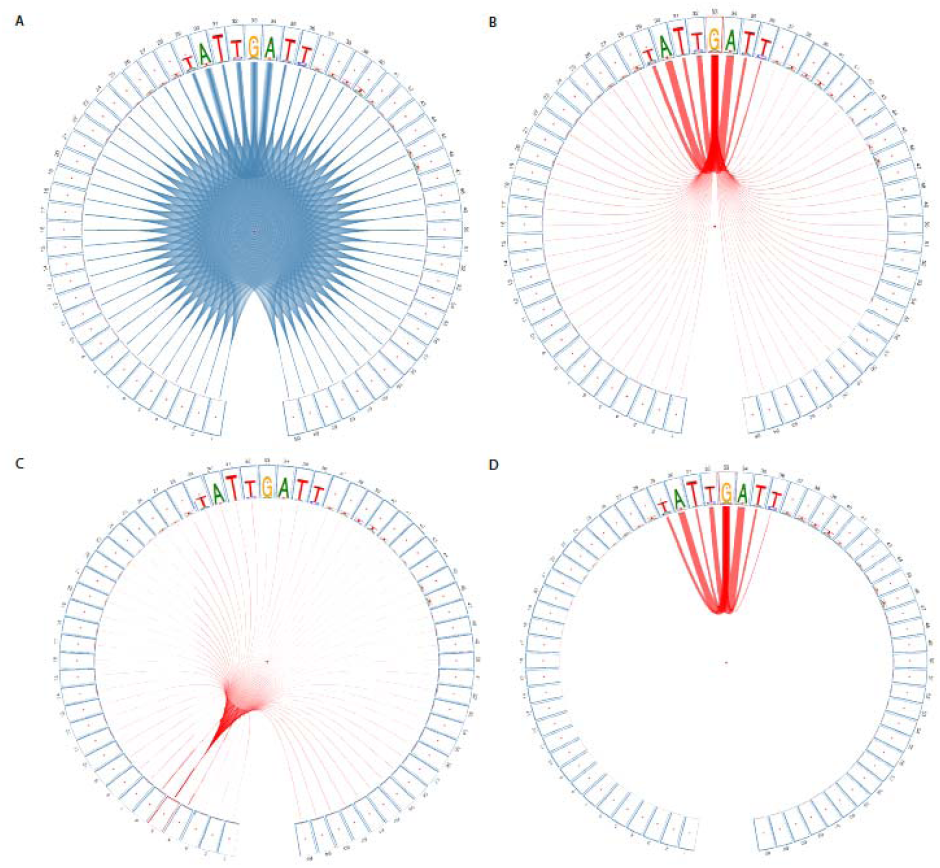
(A) Motif generated from *CircularLogo* describing the pairwise dependencies between 65 nucleotides (20 upstream nucleotides + 25 HNF6 binding sites defined from ChIP-exo data + 20 downstream nucleotides). (B) All links related to node 33. (C) All links related on node 5, representing background level dependencies. (D) Links related to node 33 after removing spurious, background links.

*CircularLogo* allows users to interactively adjust a variety of parameters and explore intra-motif dependencies and fine-tune the appearance of the final output. For example, any nucleotide in the genome has a certain level of dependencies with its immediate neighbors. We considered such dependencies as the background noise since they are not likely to be biologically meaningful. Due to genome heterogeneities, different sets of DNA motifs might have different levels of background dependency. Therefore, instead of using a fixed, hard threshold, *CircularLogo* provides users with a slider bar to interactively filter out weak background links.

### Nucleotide dependencies within HNF6 motif

HNF6 (also known as ONECUT1) is a transcription factor that regulates expression of genes involved in a variety of cellular processes. The exact protein-DNA binding boundaries of HNF6 in mouse genome were previously defined by our group[19]. A total of 5549 binding sites, each of 25 nucleotides long, were used to explore the intra-motif dependencies. Each binding site was also extended 20 nucleotides up- and downstream in order to estimate the background dependency level. Pair-wise dependencies between all 65 positions were displayed in **Fig. 1A**. As we expected, dependencies between positions within the HNF6 binding site (i.e. nucleotides within 29^th^ and 36^th^ position) were much higher than those of flanking regions (**Fig. 1B**). **Fig. 1C** indicated background links relating to node 5 (i.e. the 5^th^ position of input DNA sequence). **Fig. 1D** indicated dependencies related to node 33 within the HNF6 binding site after spurious links were removed.

### Nucleotide dependencies within tRNAs

The transfer RNA (tRNA) is involved in translating message RNA (mRNA) into the amino acid sequence. Its typical cloverleaf secondary structure is composed of D-loop, anticodon loop, variable loop and T_Ψ_C loop, as well as four base-paired stems between these loops (**Fig. 2A**). The nucleotides within stems are less conserved than those of loops, but base pairings within stems are required for structural stability. Thus we expect higher positional dependencies between nucleotides within stems than those within loops. We used *CircularLogo,* with mutual information as a measurement of dependence, to generate tRNA circular motif. After filtering out weak links (lower 33%), we observed four apparent clusters of connected links corresponding to the four stems (**Fig. 2B**). Comparing to motif logo generated from enoLOGOS (http://www.benoslab.pitt.edu/cgi-bin/enologos/enologos.cgi) using the same dataset, *CircularLogo* provided more intuitive view of intra-dependencies within the four stems (**Fig. 2C**). **Fig. 2B** also shows that nucleotides with three loops (D-loop, Anticodon loop, and T_Ψ_C loop) exhibited much higher sequence conservation than that of nucleotides located in stems, suggesting that the loops are main functional domains of tRNA. For example, D-loop is the recognition site of aminoacyl-tRNA synthetase, an enzyme involved in amino-acylation of the tRNA molecule [20,21], and T_Ψ_C loop is the recognition site of the ribosome.

**Figure 2.**
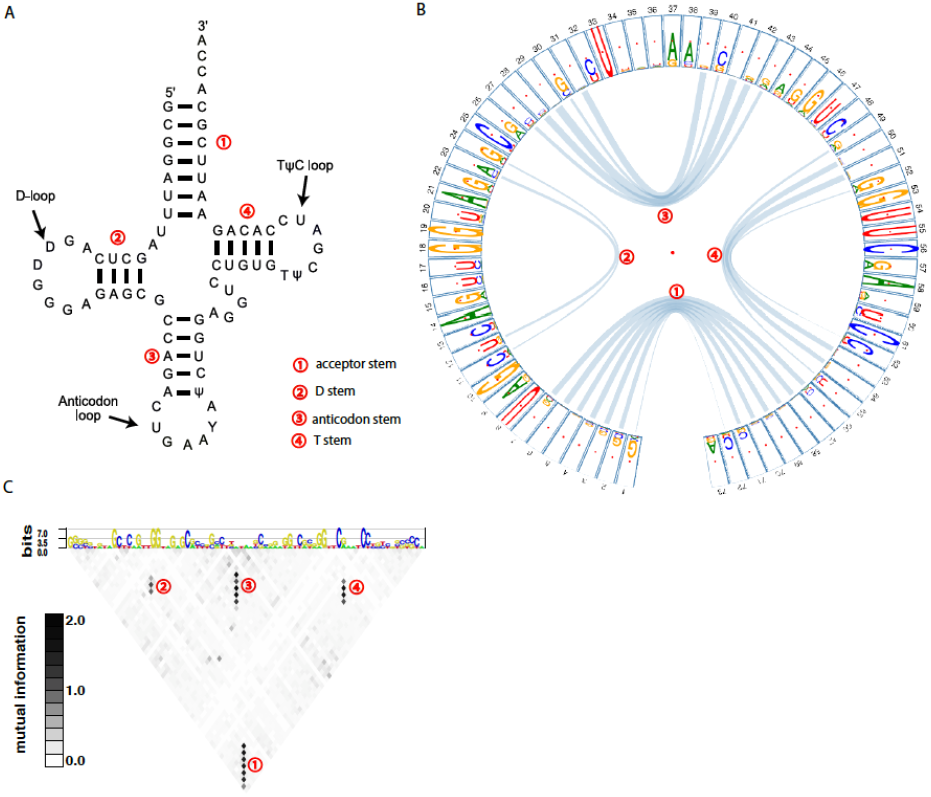
(A) The typical cloverleaf secondary structure of Phe-tRNA in yeast. (B) tRNA motif represented with the circular motif logo. The width of links indicates the strength of dependency (measured by mutual information). (C) tRNA motif logo generated from enoLOGOS using the same dataset. The labels ①, ②, ③, ④ indicate acceptor stem, D-stem, anticodon stem, and T-stem, respectively.

## Discussion

New statistical models and experimental approaches are being developed for measuring intra-motif dependency. *CircularLogo* uses a plain text, JSON-Graph formatted, file to describe DNA/RNA motifs, which enables users to generate a customized JSON-Graph file containing positional dependencies that are pre-calculated by their choice methods.

When the raw sequences were given to *CircularLogo*, it provides two approaches (χ^2^ statistic and mutual information) for measuring the positional dependency. Both of these methods, although commonly used, are biased and unable to quantify dependencies between highly conserved nucleotide stacks (e.g. invariable sites) [6,22]. This problem could be address by users providing as many sequences as possible in order to capture the low-frequent variants at those highly conserved sites. This is feasible due to genome-wide, high-throughput, screening technologies. For example, researchers usually identify tens of thousands of potential TFBSs using ChIP-seq or other similar technologies. After retrieving the potential TFBSs from ChIP-seq data, a researcher can align them using the predicted DNA motif and give the final alignment file as input for *CircularLogo*. We recommend that a FASTA input file should contain at least 20 sequences.

It is worth noting that the χ^2^ statistic and mutual information are two different measures of dependence, each suited for use under different conditions. Essentially, the χ^2^ statistic measures the co-occurrence of nucleotides of two different positions. Hence, χ^2^ method is suited for measuring dependency between two conserved (i.e. less variable) positions but it has limited power to measure dependency between two highly variable positions wherein the dinucleotide frequencies are close to background (i.e. 1/16) and the χ^2^ statistic approaches 0. In contrast, mutual information measures the reduction in uncertainty about nucleotide frequencies in one position, given some knowledge of nucleotide frequencies at another position. For a pair of highly conserved positions that are dominated by particular nucleotides, the information content of each position and the mutual information between them approaches to 0 bit. Hence, mutual information is suited for measuring dependency between two highly variable positions.

## Conclusions

Visualization is key for efficient data exploration and effective communication in scientific research. *CircularLogo* is an innovative tool offering the panorama of DNA or RNA motifs taking into consideration the intra-site dependencies. We demonstrated the utility and practicality of this tool using examples wherein *CircularLogo* was able to depict complex dependencies within motifs and reveal biomolecular structure (such as stem structures in tRNA) in an effective manner.

## List of abbreviations

BEM: the Binding Energy Model
JSON: JavaScript Object Notation
MEME: Multiple Em for Motif Elicitation
MACE: Model-based Analysis of ChIP-Exo
MI: Mutual Information
PSSM: Position-Specific Scoring Matrix
PMM: the inhomogeneous Parsimonious Markov Model
PWM: Position Weight Matrix
TFBS: Transcription Factor Binding Sites
TFFMs: Transcription Factor Flexible Model

## Declarations

### Ethics approval and consent to participate

Not applicable

### Consent for publication

Not applicable

### Availability of data and material

The datasets analyzed during the current study are available at: https://sourceforge.net/projects/circularlogo/files/test/repository

### Authors’ contributions

LW, SD and JPK conceived the study. ZY and TM implemented *CircularLogo* software and performed the analysis. MK built *CircularLogo* web server. LW, ZY, SD and JPK wrote the manuscript. All authors read and approved the final manuscript.

## Acknowledgements

Not applicable

